# Control of Oxidative Stress and Intracellular Survival in *Francisella tularensis* Live Vaccine Strain (LVS) via Acyl-CoA Synthetase

**DOI:** 10.64898/2026.05.08.723735

**Authors:** Anthony Centone, Zhuo Ma, Meenakshi Malik, Chandra Shekhar Bakshi

## Abstract

*Francisella tularensis* is a highly infectious, Gram-negative intracellular bacterium and the causative agent of tularemia, a potentially fatal disease. Owing to its low infectious dose, ease of aerosolization, high virulence, lack of an effective vaccine, and potential use as a bioterrorism agent, *F. tularensis* is classified by the CDC as a Tier 1 Category A Select Agent. Despite its clinical importance, the mechanisms underlying *F. tularensis* virulence remain incompletely understood. In this study, we generated a partial Tn5 transposon insertion mutant library in the *F. tularensis* live vaccine strain (LVS) and identified a mutant disrupted in the *FTL_0690* gene through screening under macrophage-like conditions. *FTL_0690* encodes an acyl-CoA synthetase. Characterization of both a transposon-insertion mutant and a targeted deletion mutant (Δ*FTL_0690*) revealed critical roles for this enzyme in *F. tularensis* pathobiology. Loss of *FTL_0690* increased sensitivity to oxidative stress and impaired intracellular growth within macrophages compared to wild-type *F. tularensis* LVS. Lipidomic profiling of the Δ*FTL_0690* mutant revealed disruptions in fatty acid metabolism, membrane lipid remodeling, and redox homeostasis. Altered lipid-derived and membrane-associated metabolites indicated defective phospholipid incorporation and altered membrane composition, likely contributing to oxidative stress sensitivity and reduced intramacrophage survival. Collectively, these findings demonstrate that *FTL_0690* which encodes long-chain acyl-CoA synthetase, contributes to lipid homeostasis, membrane integrity, and oxidative stress resistance of *F. tularensis*.

**Importance:** This work addresses critical gaps in our understanding of *Francisella tularensis* virulence by identifying lipid metabolism as a central determinant of intracellular survival and stress resistance. By integrating transposon mutagenesis, targeted gene deletion, and lipidomic profiling, this study provides mechanistic insight into how metabolic remodeling supports pathogenesis. Our identification and characterization of *FTL_0690* as a long-chain acyl-CoA synthetase essential for lipid homeostasis, membrane integrity, and oxidative stress resistance reveals a previously unappreciated link between fatty acid metabolism and intramacrophage survival of *F. tularensis*.

## Introduction

*Francisella tularensis* (*F. tularensis*) is a Gram-negative, intracellular bacterial pathogen and the causative agent of tularemia, a potentially lethal zoonotic disease (1). Owing to its high virulence, extremely low infectious dose, ease of aerosolization, and the associated risk to public safety, including concern over its potential use as a bioterrorism agent, *F. tularensis* is classified by the Centers for Disease Control and Prevention (CDC) as a Tier 1 Category A Select Agent (2–4). *F. tularensis* comprises four major subspecies: *F. tularensis* subsp. *tularensis*, *F. tularensis* subsp. *holarctica*, *F. tularensis* subsp. *mediasiatica*, and *F. tularensis* subsp. *novicida*. *F. tularensis* subsp. *tularensis* (Type A) is the most virulent subspecies and includes the highly pathogenic SchuS4 strain; it is endemic to North America (5). In contrast, *F. tularensis* subsp. *holarctica* (Type B) is less virulent but is nonetheless capable of causing severe human disease (6). The widely studied Live Vaccine Strain (LVS) is a derivative of *F. tularensis* subspecies *holarctica* (7). The remaining subspecies, *F. tularensis* subsp. *novicida* exhibits limited pathogenicity in healthy humans but can cause disease in immunocompromised individuals (8). *F. tularensis* subsp. *mediasiatica* is primarily of ecological relevance and is not associated with any human disease (9).

Tularemia can manifest in several distinct clinical forms depending on the route of transmission. The most common presentations include cutaneous, ulceroglandular, oculoglandular, lymphadenopathic, typhoidal, and pneumonic disease. Transmission through bites or skin contact often results in localized ulcers at the site of entry, accompanied by regional lymphadenopathy, whereas transmission via the oral or ocular mucosa commonly presents with cervical or preauricular lymph node enlargement (10–12). Pneumonic tularemia, which may arise from direct inhalation of *F. tularensis* or through hematogenous dissemination from other sites of infection, is the most severe form and is associated with mortality rates approaching 60% if left untreated (12, 13).

Acyl-CoA synthetases catalyze the ATP-dependent activation of free fatty acids through thioesterification to coenzyme A, generating acyl-CoA intermediates that are essential for bacterial lipid metabolism. This conserved two-step reaction, which proceeds via a fatty acyl-adenylate intermediate, is required for fatty acid β-oxidation, membrane phospholipid and lipopolysaccharide biosynthesis, lipid remodeling, protein acylation, and the synthesis of complex virulence-associated lipids (14). As such, acyl-CoA synthetases occupy a central metabolic position linking nutrient acquisition to membrane homeostasis, stress adaptation, and pathogenic fitness. In *Escherichia coli*, the fatty acyl-CoA synthetase FadD couples long-chain fatty acid uptake with intracellular activation, enabling efficient assimilation of exogenous fatty acids and recycling of endogenous lipids released during membrane turnover (15, 16). In pathogenic bacteria, expanded and specialized repertoires of acyl-CoA synthetases, such as those found in *Mycobacterium tuberculosis* and *Pseudomonas aeruginosa*, facilitate exploitation of host lipid pools, resistance to oxidative and acidic stress, lipid remodeling, and regulation of virulence (17–19).

Although less well characterized at the enzymatic level, accumulating evidence indicates that *F. tularensis* similarly exploits host lipid reserves, including triacylglycerols stored in lipid droplets, by inducing host lipolysis through AMP-activated protein kinase signaling, thereby liberating free fatty acids that support bacterial replication (20). Genome-wide metabolic analyses further reveal that *F. tularensis* encodes functional pathways for fatty acid β-oxidation and acyl-CoA synthesis, consistent with active fatty acid utilization during intracellular growth (21–24). Given the critical dependence of *F. tularensis* on membrane integrity and oxidative stress resistance, acyl-CoA synthetase-mediated lipid activation represents a key yet underexplored determinant of bacterial fitness and virulence. Our work provides the first direct characterization of the role of an acyl-CoA synthetase in *F. tularensis*.

## Materials and Methods

### Bacterial Strains, stock preparation, and maintenance

The *Francisella tularensis* live vaccine strain (LVS) used in this study was obtained from BEI Resources (Manassas, VA, USA). *F. tularensis* LVS was routinely cultured on Mueller-Hinton (MH) chocolate agar plates (Hardy Diagnostics, Santa Maria, CA, USA). Liquid cultures were grown in Mueller-Hinton broth (MHB; BD Bacto, Franklin Lakes, NJ, USA) supplemented with 0.25% ferric pyrophosphate and 2% IsoVitalex enrichment (BD Bacto). For preparation of bacterial stocks, *F. tularensis* LVS was grown on MH chocolate agar plates to mid-logarithmic phase, harvested, and resuspended in MHB to an optical density at 600 nm greater than 1.0. Aliquots were flash-frozen in liquid nitrogen and stored at −80°C. Transposon-insertion mutants and a targeted gene-deletion mutant of the acyl-CoA synthetase gene (Δ*FTL_0690*), along with its trans-complemented strain (Δ*FTL_0690* + p*FTL_0690*), were generated in this study. Transposon-insertion mutants were maintained on MH-chocolate agar supplemented with kanamycin (25 µg mL⁻¹; MilliporeSigma), while the trans-complemented strain was cultured on MH-chocolate agar containing hygromycin B (200 µg mL⁻¹; MilliporeSigma). Stock cultures of all mutant strains were prepared as described for wild-type *F. tularensis* LVS. *Escherichia coli* strain NEB 5-alpha (New England Biolabs, Ipswich, MA, USA; Cat. No. C2987H) was used as the cloning host for all recombinant DNA procedures. For stock preparation, *E. coli* cultures were grown to mid-logarithmic phase on Luria-Bertani (LB) agar, resuspended in LB broth supplemented with 15% (v/v) glycerol, and stored at −80°C.

### Cell Culture

All mammalian cell culture work was performed using RAW264.7 murine macrophages (ATCC TIB-71; American Type Culture Collection, Manassas, VA, USA). Cells were maintained in Dulbecco’s Modified Eagle Medium (DMEM; high glucose, with pyruvate; Gibco, Thermo Fisher Scientific) supplemented with 10% fetal bovine serum (FBS; Gibco, Thermo Fisher Scientific). Cultures were incubated at 37°C in a humidified atmosphere containing 5% CO₂.

### Generation of transposon-insertion mutant library

Random transposon mutagenesis was performed using the EZ-Tn5™ *KAN-2* Insertion System (Epicenter, Madison, WI, USA) according to the manufacturer’s instructions. The Tn5 transposon insert was excised from the pMOD-2 plasmid by restriction digestion, separated by agarose gel electrophoresis, and purified before use. Purified transposon DNA was then mixed with transposase and introduced into electrocompetent *F. tularensis* cells by electroporation as previously described (25, 26). Following transformation, bacteria were plated on MH-chocolate agar supplemented with kanamycin (25 µg mL⁻¹) to select for transposon insertion mutants. Individual kanamycin-resistant colonies were picked, resuspended in sterile MH broth in 96-well plates, and incubated at 37°C for 24 hours. Glycerol was subsequently added to each well to a final concentration of 15%, and the resulting *F. tularensis* Tn5 mutant library was stored at −80°C until further use.

### Screening of transposon mutants for sensitivity to pH, nitrosative and oxidative stress

Susceptibility of wild-type and transposon mutants to various stressors, such as pH, oxidants, and detergents. For pH sensitivity, transposon-insertion mutants were cultured in BHI broth adjusted to pH 6.8 for 24 hours (OD₆₀₀ = ∼ 0.2). Following this, the cultures were diluted 10-fold in BHI medium adjusted to pH 5.6 or 6.8 and incubated in a shaker incubator at 37°C for 3 hours. Each culture was spotted onto MH-chocolate agar plates to assess viability and identify pH-sensitive mutants.

For oxidant sensitivity assays, cultures of wild-type *F. tularensis* LVS and transposon insertion mutants grown on MH-chocolate agar plates were harvested and resuspended in Brain-Heart Infusion broth to an OD_600_ of 0.2. Cultures were exposed to 0.84 mM sodium nitroprusside or 1.2 mM pyrogallol and incubated at 37°C with shaking for 3 hours. Following exposure, aliquots were spotted onto MH-chocolate agar plates to assess viability. Untreated cultures served as controls. The gene-deletion mutant and the transcomplemented strains were also tested for oxidant sensitivity and sodium dodecyl sulfate (SDS) using the protocols described above.

### Identification of transposon insertion sites

Transposon insertion sites were mapped using a custom reference-guided contig-screening workflow developed to address computational limitations. Whole-genome sequencing of individual mutant strains was performed by Azenta Genewiz (South Plainfield, NJ, USA) using Illumina paired-end 150-bp sequencing. Sequencing reads were aligned to the *Francisella tularensis* LVS reference genome (NCBI accession no. AM233362.1) using a reference-guided assembly approach, which assembled aligned sequences while segregating unaligned contigs into a separate dataset. Unaligned contigs were screened for the presence of the EZ-Tn5 *KAN-2* mosaic end sequence using Geneious Prime. Contigs containing transposon sequence were analyzed to identify adjacent genomic flanking regions, which were aligned to the reference genome to determine the precise chromosomal insertion sites. Insertion site assignments were validated by transposon-flanking PCR, with successful amplification confirming transposon integration. The primer sequences used are detailed in **Table 1**.

**Table 1.**
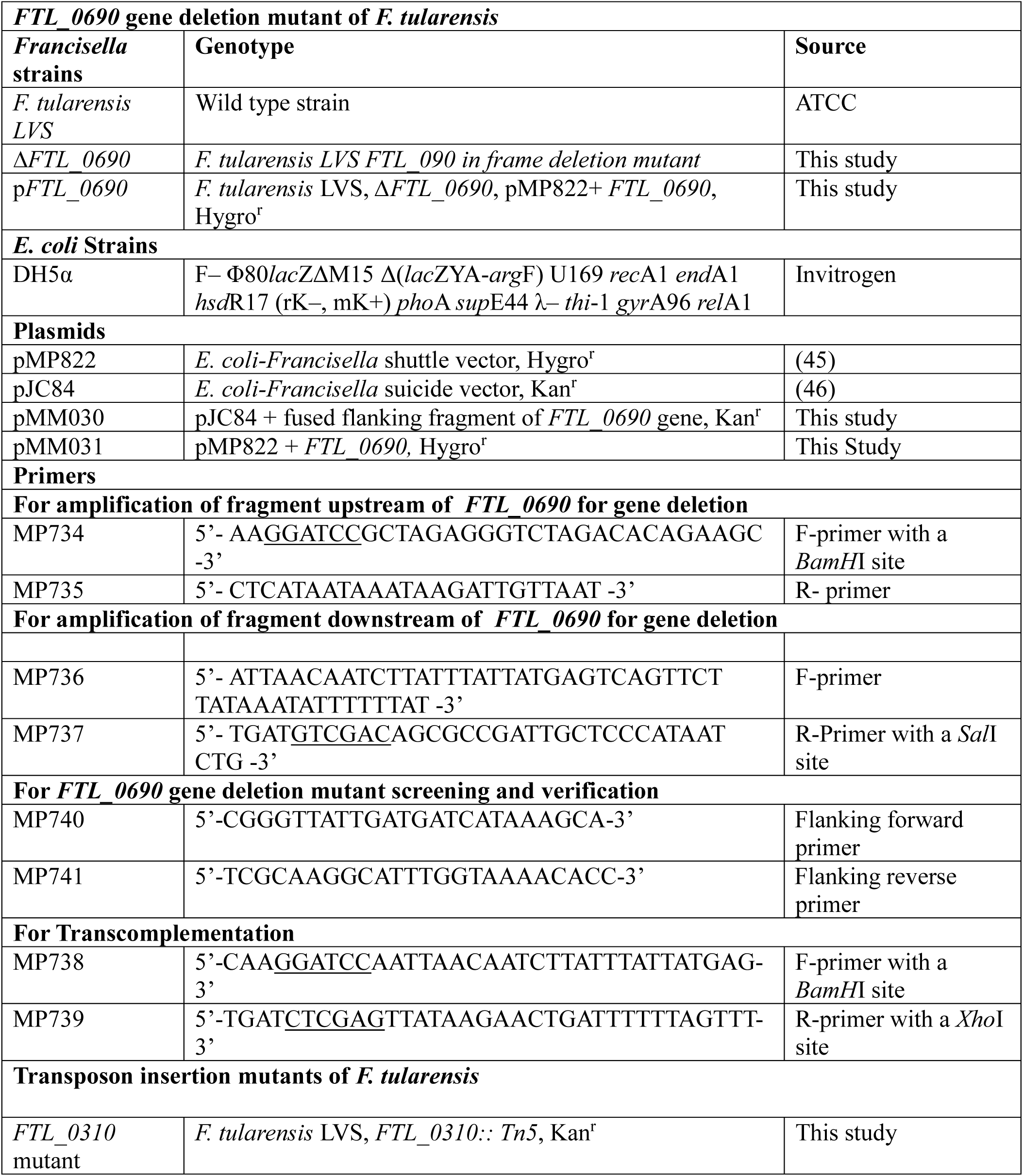
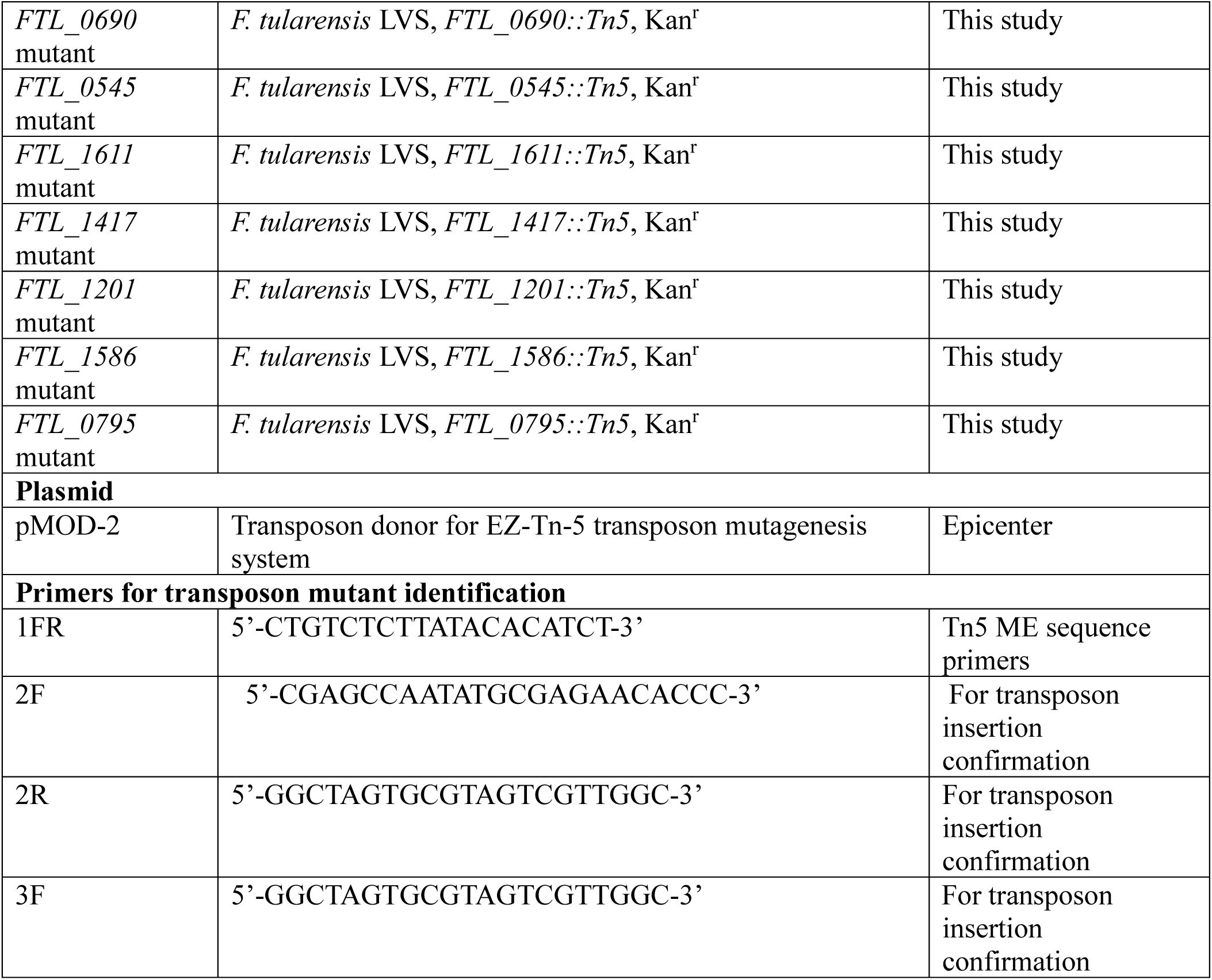
List of *Francisella tularensis* LVS strains developed, plasmids and primers used in the study.

### Generation of *ΔFTL_0690* and the transcomplemented strains of *F. tularensis* LVS

An in-frame deletion mutant of the *F. tularensis* acyl-CoA synthetase gene (*FTL_0690*) was generated using allelic exchange as described earlier (27, 28). Briefly, fragments corresponding to regions upstream and downstream of the *FTL_0690* coding sequence were amplified and fused to generate a deletion construct lacking the majority of the *FTL_0690* open reading frame. The resulting deletion allele was cloned into a suicide vector pJC84. The resulting plasmid pMM030 was transformed into *F. tularensis* LVS by electroporation. Double-crossover recombinants were selected using Kanamycin selection and 8% sucrose counter-selection. The *FTL_0690* gene deletion was confirmed by PCR using the gene-specific flanking primers MP740 and MP741. For transcomplementation, the full-length *FTL_0690* gene was amplified from *F. tularensis* LVS genomic DNA and cloned into a pMP822 *E. coli-Francisella* shuttle vector. The resulting plasmid (pMM031) was introduced into the Δ*FTL_0690* mutant strain by electroporation to generate the transcomplemented strain (*ΔFTL_0690* + p*FTL_0690*). The transcomplemented strains were selected on MH chocolate agar plates containing hygromycin B and maintained under antibiotic selection. The primer sequences used are detailed in **Table 1**.

### Macrophage invasion assay

The macrophage invasion assays were conducted as described earlier (25, 29, 30). Briefly, RAW264.7 murine macrophages were seeded in 6-well tissue culture plates at a density of 1 × 10⁶ cells per well and incubated for 16 hours before infection. Bacterial suspensions were prepared in complete cell culture medium and adjusted to 10 or 100 multiplicity of infection (MOI). Infections were initiated by adding bacteria directly to each well, followed by centrifugation at 1,000 × g for 10 min at 4°C to promote bacterial contact. Plates were then incubated at 37°C with 5% CO₂ for 2 hours to allow internalization. Following infection, cells were washed with PBS and incubated with medium containing gentamicin (50 µg mL⁻¹) for 1 hour to eliminate extracellular bacteria. After antibiotic treatment, cells were washed and maintained in antibiotic-free medium for the remainder of the experiment. At 4 and 24 hours post-infection, cells were lysed in DMEM containing 0.1% sodium deoxycholate, and the lysates were diluted 10-fold and plated on MH chocolate agar plates to enumerate intracellular bacteria. Results were expressed as colony-forming units (CFU) per milliliter.

### Untargeted metabolomics by mass spectrometry

Untargeted metabolomics analysis was performed by Creative Proteomics (Shirley, NY, USA). Wild-type and Δ*FTL_0690 F. tularensis* strains were grown to mid-logarithmic phase on chocolate agar plates. Bacterial growth was harvested, resuspended in PBS, and pelleted by centrifugation at 1,000 × g for 10 min at 4°C. Approximately 2 × 10¹⁰ cells (OD₆₀₀ = 4.0) were collected per strain in biological triplicate. Pellets were flash-frozen in liquid nitrogen and shipped overnight to Creative Proteomics on dry ice. Metabolite extraction, liquid chromatography-tandem mass spectrometry (LC-MS/MS) analysis, and downstream bioinformatic processing were conducted by Creative Proteomics. Metabolites were separated by ultra-high-performance liquid chromatography (UHPLC) and detected in both positive (ESI⁺) and negative (ESI⁻) ionization modes. Raw spectral data were processed for peak detection, alignment, normalization, and compound identification using internal and public metabolite databases. Multivariate and univariate statistical analyses, including principal component analysis (PCA), variable importance in projection (VIP) scoring, and volcano plot analysis, were performed to identify metabolites significantly altered between wild-type and Δ*FTL_0690* strains.

### Mice Experiments

All animal studies were conducted in accordance with protocols approved by the New York Medical College Institutional Animal Care and Use Committee (IACUC protocol no. 19295). Female C57BL/6J mice (6-8 weeks old; Charles River Laboratories) were acclimated for 2-7 days prior to experimentation. Mice were anesthetized by intraperitoneal injection of a ketamine/xylazine cocktail before infection. Wild-type and Δ*FTL_0690 F. tularensis* strains were prepared in PBS, and mice were inoculated intranasally with 2 × 10⁴, 2 × 10⁵, or 2 × 10⁶ CFUs. Intranasal inoculations were performed in a total volume of 20 µL (10 µL per nostril). Mice were monitored daily for weight changes until all animals in a group had either succumbed to infection or fully recovered. Humane endpoints were necessary in severely moribund mice. At the end of the experiment, surviving mice were euthanized by isoflurane anesthesia followed by cervical dislocation. Inoculum doses were confirmed by serial dilution and plating for CFU enumeration immediately following inoculation.

### Statistical analysis

Experiments were performed with three technical replicates per condition unless otherwise specified. All statistical analyses were performed using GraphPad Prism or Microsoft Excel. Data are presented as mean ± standard error of the mean (SEM) unless otherwise indicated. Statistical significance was assessed using unpaired two-tailed Student’s *t*-tests for pairwise comparisons and one-way or two-way analysis of variance (ANOVA) for multiple comparisons, as appropriate. A *p*-value < 0.05 was considered statistically significant.

## Results

### Screening of transposon insertion mutants led to the identification of genes responsible for pH resistance

Transposon mutagenesis generated a library of 765 individual *F. tularensis* mutant colonies, which covered approximately 50% of the *Francisella* genome. This library was screened under conditions designed to model key intramacrophage stressors encountered during host infection: acidic pH of 5.6 to mimic phagosomal acidification and oxidative stress. Nineteen mutants were identified that showed increased sensitivity to pH 5.6 (Fig. 1A). Eleven of these mutants were excluded from further analysis because they also exhibited growth defects under normal pH conditions on MH-chocolate agar plates, indicating nonspecific attenuation. The remaining 8 mutants were sequenced to identify the gene interrupted by the transposon insertion. Whole-genome sequencing identified disrupted loci with putative functional roles in *F. tularensis*. The *FTL_0310*, which encodes the E2 component of the pyruvate dehydrogenase complex, functions in acetyl-CoA generation and connects central carbon metabolism to TCA-cycle-associated stress defense. *FTL_0545*, encoding phosphatidylcholine synthase, functions in phospholipid remodeling and participates in phosphatidylethanolamine synthesis and modulation of host inflammatory lipid responses during macrophage infection. *FTL_1611*, a group 2 glycosyltransferase, functions in polysaccharide biosynthesis and contributes to cell wall and lipopolysaccharide biogenesis. *FTL_1417*, encoding a major facilitator superfamily transporter, functions in membrane-associated solute transport and is located immediately upstream of capB, a tularemia vaccine-associated locus. *FTL_1200*, a p47K family protein, functions in cobalamin (vitamin B12) biosynthesis and belongs to a regulatory network controlled by the PmrA response regulator required for intramacrophage growth. *FTL_1585*, encoding a RecJ-like single-stranded DNA-specific exonuclease, functions in DNA repair and homologous recombination. One transposon insertion mapped to an intergenic region between *FTL_0794* and *FTL_0795*, adjacent to an adenylate kinase locus, and did not disrupt a characterized coding sequence. *FTL_0690*, encoding a long-chain fatty-acid-CoA ligase (acyl-CoA synthetase), functions in fatty acid activation for β-oxidation and membrane phospholipid incorporation, was identified (Fig 1B).

**FIGURE 1.**
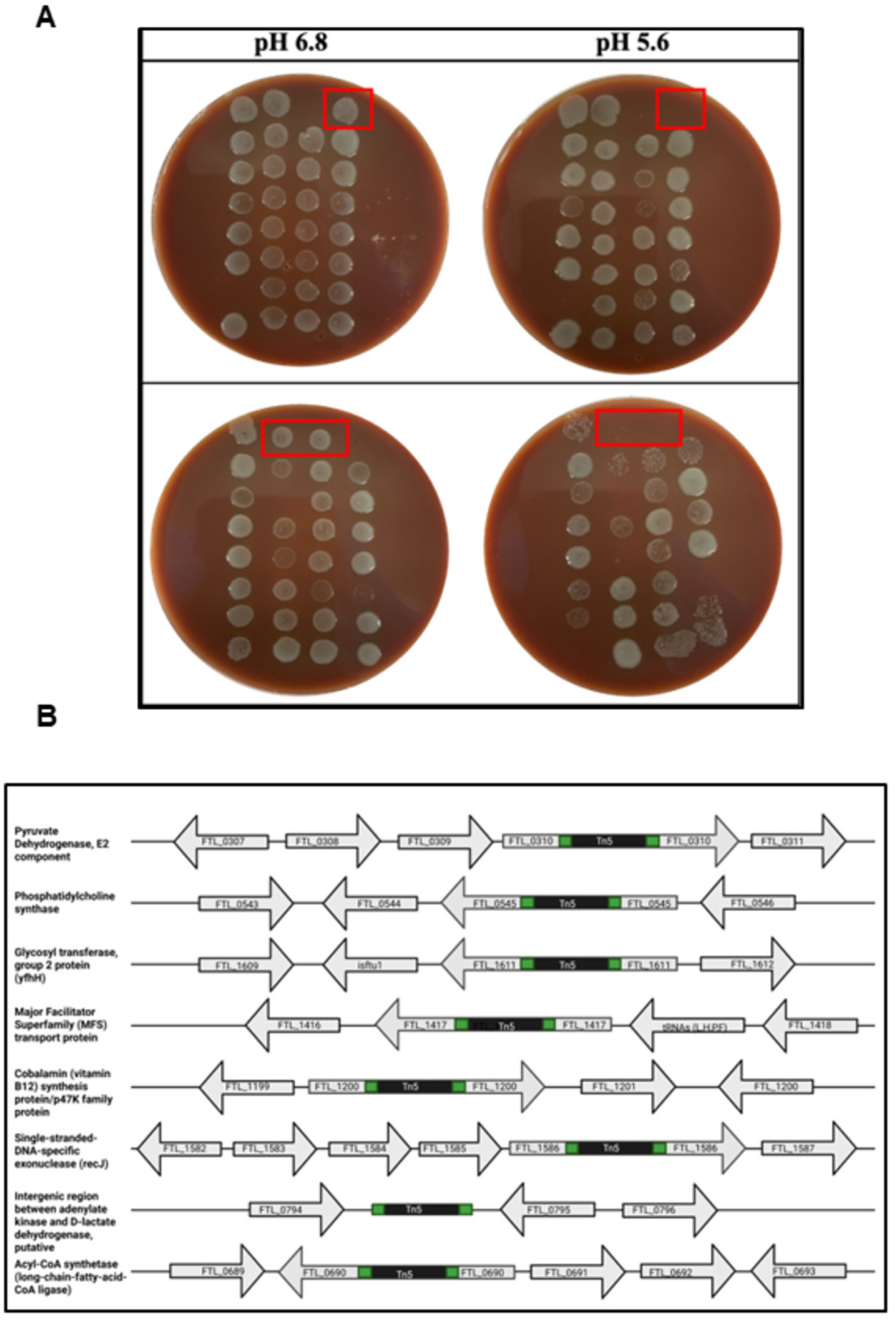
Identification of pH-sensitive transposon insertion mutants of *F. tularensis* LVS and mapping of disrupted genes. **(A)** Transposon-insertion mutants were cultured in BHI broth adjusted to pH 6.8 for 24 hours (OD₆₀₀ = ∼ 0.2). Following this, the cultures were diluted 10-fold in BHI medium adjusted to pH 5.6 or 6.8 and incubated in a shaker incubator at 37°C for 3 hours. Each culture was spotted onto MH-chocolate agar plates to assess viability and identify pH-sensitive mutants. Mutants that failed to grow after exposure to pH 5.6 were identified as pH-sensitive (red boxes). Representative images from the screening of 765 transposon-insertion mutants are shown. **(B)** The identified pH-sensitive mutants were subjected to whole-genome sequencing to map the genes interrupted by transposon insertions. The black boxes indicate the site(s) of transposon insertion.

### Characterization of transposon insertion mutants for their sensitivity to nitrosative, oxidative stresses, and intramacrophage survival

To evaluate the contribution of genes identified by transposon mutagenesis to nitrosative stress resistance, wild-type *F. tularensis* LVS and selected transposon insertion mutants were assessed for in the presence of the nitric oxide donor sodium nitroprusside (0.84 mM). Wild-type *F. tularensis* LVS exhibited comparable growth under untreated conditions and in the presence of sodium nitroprusside, indicating intrinsic resistance to nitrosative stress. Among the mutants tested, the *Tn5:FTL_0545,* followed by the *Tn5:FTL_0690, Tn5:FTL_1585* and *Tn5:FTL_0794-FTL_0795* intergenic transposon mutant displayed a marked growth defect in the presence of sodium nitroprusside compared to untreated conditions. In contrast, other mutants, including *Tn5:FTL_0310, Tn5:FTL_1417* and *Tn5:FTL_1200* mutant, showed growth patterns similar to wild-type *F. tularensis* LVS under nitrosative stress (Fig. 2A). These results demonstrate that disruption of *FTL_0545*, which encodes phosphatidylcholine synthase, and *FTL_0690,* which encodes acetyl-CoA synthetase, specifically compromises resistance to nitrosative stress. In contrast, disruption of the other gene loci does not markedly affect survival under nitric oxide exposure.

**FIGURE 2.**
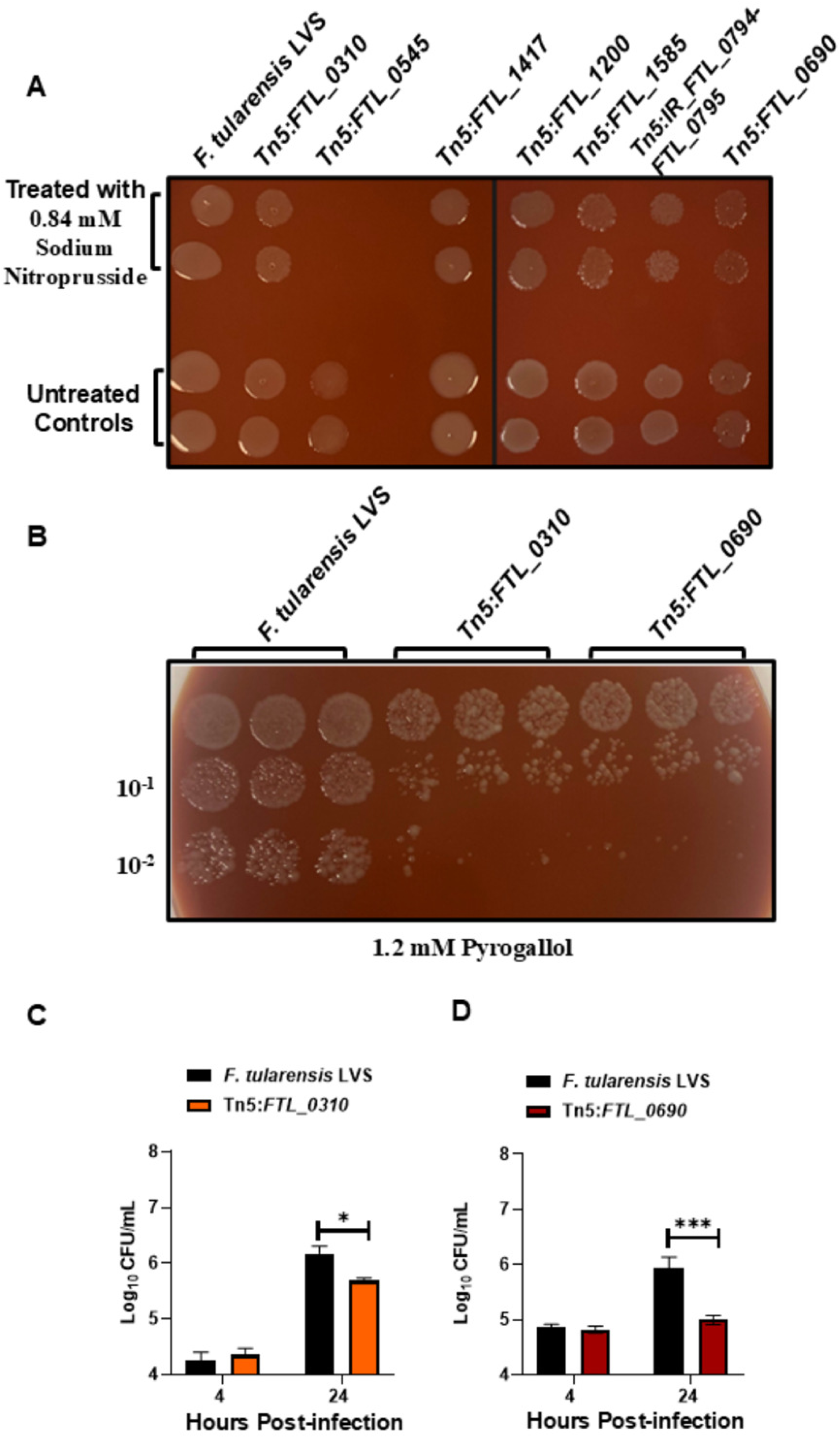
Characterization of transposon mutants of *F. tularensis* LVS for their sensitivity to nitrosative and oxidative stress, and intramacrophage survival. **(A)** Cultures of wild-type *F. tularensis* LVS and indicated transposon insertion mutants grown on MH-chocolate agar plates were harvested and resuspended in Brain-Heart Infusion broth to an OD_600_ of 0.2. Cultures were exposed to 0.84 mM sodium nitroprusside and incubated at 37°C with shaking for 3 hours. Following exposure, aliquots were spotted onto MH-chocolate agar plates to assess viability. Untreated cultures served as controls. Each spot represents an independent technical replicate (n = 2). **(B)** Cultures of wild-type *F. tularensis* LVS and the indicated transposon mutants were grown as in A, exposed to 1.2 mM pyragallol, and incubated at 37°C with shaking for 3 hours. Following exposure, aliquots were spotted onto MH-chocolate agar plates to assess viability. Untreated cultures served as controls. Each spot represents an independent technical replicate (n = 3). **(C, D)** RAW264.7 macrophages were infected with (C) *Tn5:FTL_0310* and **(D)** Tn5:*FTL_0690* mutants at a multiplicity of infection (MOI) of 10. Wild-type *F. tularensis* LVS was used as a control. Intracellular bacteria were quantified by lysing infected macrophages at 4 and 24 hours post-infection. Bacterial counts are expressed as Log₁₀ CFU/ml. Data represent the mean ± SEM from three independent experiments. Statistical significance was determined using one-way ANOVA. \**P* ≤ 0.05, \*\*\**P* ≤ 0.001.

To further assess the role of genes identified by transposon mutagenesis in resistance to oxidative stress, wild-type *F. tularensis* LVS and *Tn5:FTL_0130* and *Tn5:FTL_0690* mutants were examined for their sensitivity to superoxide-generating compound pyrogallol (1.2 mM). The *Tn5:FTL_0310* mutant exhibited a pronounced reduction in growth in the presence of pyrogallol, consistent with enhanced sensitivity to oxidative stress. Similarly, the *Tn5*:*FTL_0690* mutant showed impaired growth relative to wild-type *F. tularensis* LVS (Fig. 2B). Under untreated conditions, both mutants grew comparably to wild-type *F. tularensis* LVS (not shown), indicating that the observed defects were specific to oxidative stress exposure. These results indicate that disruption of *FTL_0310*, which encodes the E2 component of the pyruvate dehydrogenase complex, and *FTL_0690*, which encodes an acyl-CoA synthetase, increases sensitivity to oxidative stress relative to wild-type *F. tularensis* LVS. The oxidative stress sensitivity, especially of the *Tn5:FTL_0690* mutant, highlights the importance of fatty-acid pathways in supporting bacterial survival under reactive nitrogen/oxygen species-mediated stress.

To evaluate the role of identified genes in macrophage invasion and intracellular survival, RAW264.7 murine macrophages were infected with wild-type *F. tularensis* LVS or *Tn5:FTL_0310* and *Tn5:FTL_0690* mutants at 10 MOI. Intracellular bacteria were quantified at 4 and 24 hours post-infection following gentamicin protection. At 4 hours post-infection, intracellular bacterial burdens for both the *Tn5:FTL_0310* and *Tn5:FTL_0690* mutants were comparable to wild-type *F. tularensis* LVS, with no statistically significant differences observed, indicating that disruption of either gene did not impair macrophage entry or early intracellular survival. By 24 hours post-infection, wild-type *F. tularensis* LVS replicated efficiently within macrophages, resulting in a significant increase in intracellular CFU. In contrast, both *Tn5:FTL_0310* and *Tn5:FTL_0690* mutants exhibited significantly lower intracellular bacterial burdens than the wild-type control. The *Tn5:FTL_0310* mutant displayed a clear defect in intracellular replication, while the *Tn5:FTL_0690* mutant showed a pronounced reduction in CFU, demonstrating impaired persistence or replication within macrophages (Fig. 2C and D). Together, these results show that *FTL_0310* and *FTL_0690* are not required for bacterial entry into macrophages but are essential for sustained intracellular replication and survival during later stages of infection. The results, especially for the *Tn5:FTL_0690* mutant, highlight the importance of fatty acid metabolism in the nitrosative and oxidative stress response and, in particular, the intracellular growth of *F. tularensis* within macrophages.

### Genomic context of *FTL_0690* and confirmation of deletion and complementation

Analysis of the *FTL_0690* genomic locus showed that *FTL_0690* is positioned downstream of *FTL_0692* and *FTL_0691* genes. All these genes are oriented in the same transcriptional direction, consistent with operon-level organization. *FTL_0691* and *FTL_0692* encode a proton-dependent oligopeptide transport protein and a hypothetical conserved protein, respectively. This genomic organization suggests that *FTL_0691* and *FTL_0692* function as accessory or regulatory components involved in fatty acid activation and adaptation to metabolic stress. The gene upstream of *FTL_0690* is characterized as an oxidative stress response regulator (OsrR) belonging to the AraC/XylS family of transcriptional regulators (31) *FTL_0690* encodes a 563 amino acid long-chain-fatty acid CoA-ligase (acyl-CoA-Synthetase), similar to FadD of *E. coli*.

PCR analysis confirmed the deletion of the *FTL_0690* gene. Amplification of the *FTL_0690* locus from wild-type *F. tularensis* LVS yielded a product of the expected size (∼1922 base pairs). In contrast, the Δ*FTL_0690* mutant produced a truncated amplicon of approximately 268 base pairs, consistent with deletion of the *FTL_0690* coding region. Introduction of the transcomplementation plasmid (*pFTL_0690*) into the Δ*FTL_0690* mutant resulted in a full-length amplicon, confirming successful transcomplementation (Fig. 3B).

**FIGURE 3.**
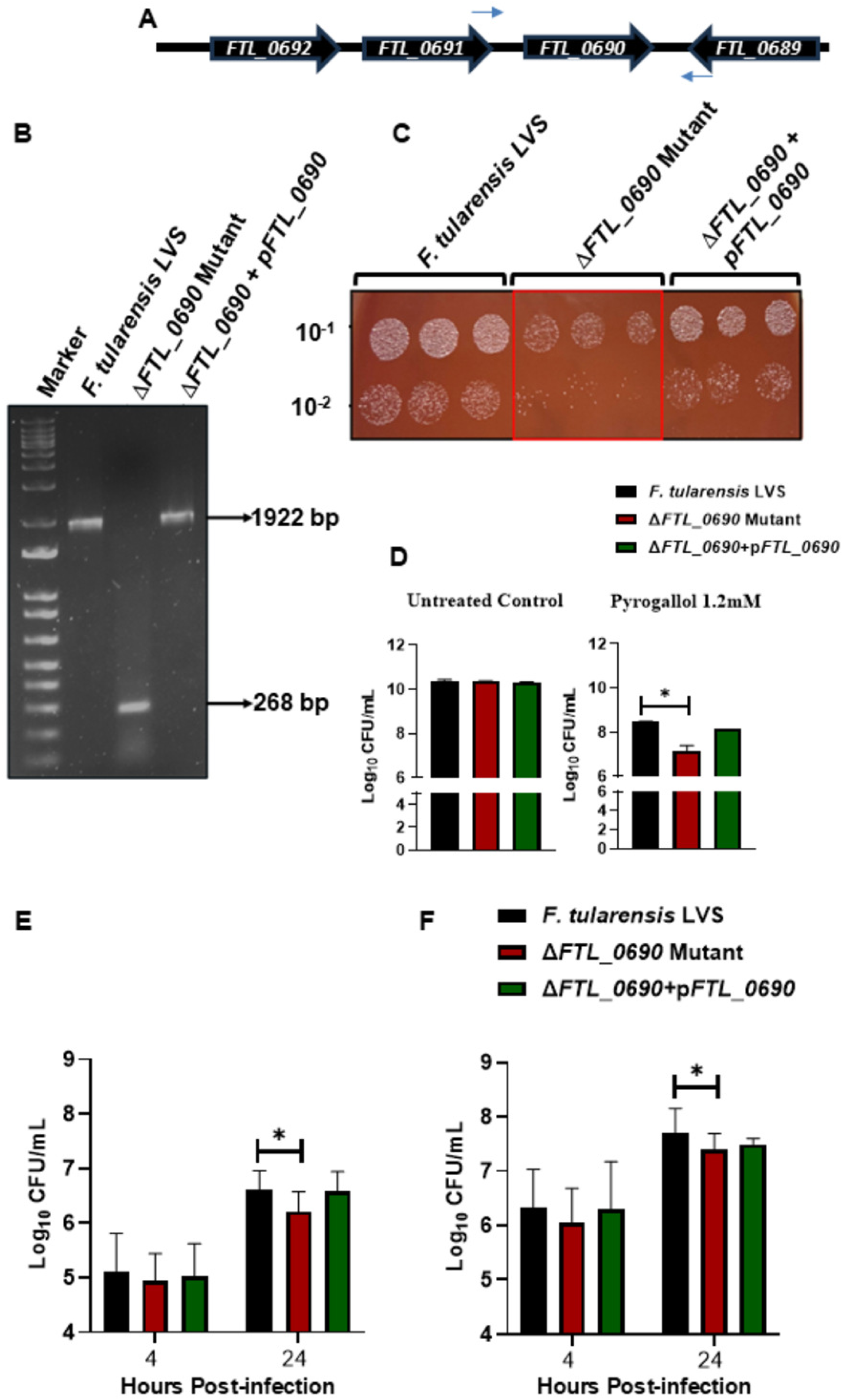
Generation and characterization of the *FTL_0690* gene deletion mutant of *F. tularensis* LVS. **(A)** Genomic organization of the *FTL_0690* gene encoding acyl-CoA-synthetase of *F. tularensis* LVS. The blue arrows indicate the location of the primers (MP740 and MP741) used to confirm *FTL_0690* gene deletion. **(B)** Confirmation of gene deletion (Δ*FTL_0690*) and transcomplemented (Δ*FTL_0690* + p*FTL_0690*) strains of *F. tularensis* LVS. **(C)** Cultures of the indicated strains were grown on MH-chocolate agar plates, harvested, and resuspended in Brain-Heart Infusion broth to an OD_600_ of 0.2. Cultures were exposed to 1.2 mM pyragallol and incubated at 37°C with shaking for 3 hours. Following exposure, aliquots were spotted onto MH-chocolate agar plates to assess viability, or **(D)** were serially diluted and plated to enumerate colony-forming units (CFUs). Untreated cultures served as controls. Each spot in C represents an independent technical replicate (n = 3). **(E, F)** RAW264.7 macrophages were infected with the indicated strains at a multiplicity of infection (MOI) of 10 **(E)** and 100 **(F)**. Intracellular bacteria were quantified by lysing infected macrophages at 4 and 24 hours post-infection. Bacterial counts are expressed as Log₁₀ CFU/ml. Data in D and E represent the mean ± SEM from three independent experiments. Statistical significance was determined using one-way ANOVA. \**P* ≤ 0.05.

### Confirmation of *FTL_0690*-dependent oxidative stress resistance and intramacrophage survival

To validate that the oxidant sensitivity observed in the *Tn5:FTL_0690* mutant was directly attributable to the disruption of the *FTL_0690* gene and not to polar effects, the Δ*FTL_0690* mutant and the transcomplemented strain were tested for their sensitivity to oxidative stress induced by the superoxide-generating compound pyrogallol. Wild-type *F. tularensis* LVS exhibited robust growth, whereas the Δ*FTL_0690* mutant displayed a pronounced growth defect in the presence of 1.2 mM pyrogallol. The Δ*FTL_0690* + p*FTL_0690* strain showed growth comparable to wild-type *F. tularensis* LVS, confirming functional transcomplementation (Fig. 3C). Quantitative analysis of bacterial survival under untreated conditions showed no significant differences in CFU among wild-type *F. tularensis* LVS, the Δ*FTL_0690* mutant, and the transcomplemented strain, indicating that deletion of *FTL_0690* did not affect basal growth. In contrast, exposure to 1.2 mM pyrogallol resulted in a significant reduction in the viability of the Δ*FTL_0690* mutant relative to wild-type *F. tularensis* LVS. This oxidative stress sensitivity was reversed in the transcomplemented strain, similar to the wild-type bacteria (Fig. 3D).

To further confirm that the macrophage replication defect observed in the *Tn5:FTL_0690* transposon mutant was specifically due to the disruption of *FTL_0690* and not polar effects, RAW 264.7 macrophages were infected with wild-type *F. tularensis* LVS, the Δ*FTL_0690* deletion mutant, and the complemented strain (Δ*FTL_0690* + p*FTL_0690*) at MOI of 10 and 100. Intracellular bacteria were quantified at 4 and 24 hours post-infection. At 4 hours post-infection, intracellular CFU counts for the Δ*FTL_0690* mutant remained similar to wild-type *F. tularensis* LVS and the transcomplemented strain at both MOIs. By 24 hours post-infection, wild-type *F. tularensis* LVS demonstrated intracellular replication at both MOIs. In contrast, the Δ*FTL_0690* mutant exhibited significantly reduced intracellular growth, with lower CFU counts relative to wild-type bacteria (Fig. 3E and F). Together, these data confirm that the loss of acyl-CoA synthetase encoded by the *FTL_0690* gene enhances sensitivity to superoxide-mediated oxidative stress and impairs survival of *F. tularensis* in macrophages. The concordance between the deletion mutant and the transposon mutant phenotypes of *FTL_0690*, coupled with complementation, establishes that these defects are specifically attributable to the loss of *FTL_0690* and not to secondary polar effects on neighboring genes, especially the *orsR* (*FTL_0689*) gene, which has also been shown to play a role in oxidative stress resistance and intracellular survival of *F. tularensis* (31).

### Metabolic alterations associated with the deletion of the *FTL_0690* gene

Untargeted metabolomic profiling was performed to identify metabolic changes associated with the deletion of *FTL_0690* in *F. tularensis* LVS. Analysis was conducted using UPLC-MS under both positive (ESI⁺) and negative (ESI⁻) electrospray ionization modes. Principal component analysis (PCA) revealed clear separation between wild-type *F. tularensis* LVS and the Δ*FTL_0690* mutant in both ionization modes, indicating substantial global differences in metabolic profiles between the two strains. Tight clustering of technical replicates within each group demonstrated high reproducibility of the metabolomic datasets (Fig. 4A).

**FIGURE 4.**
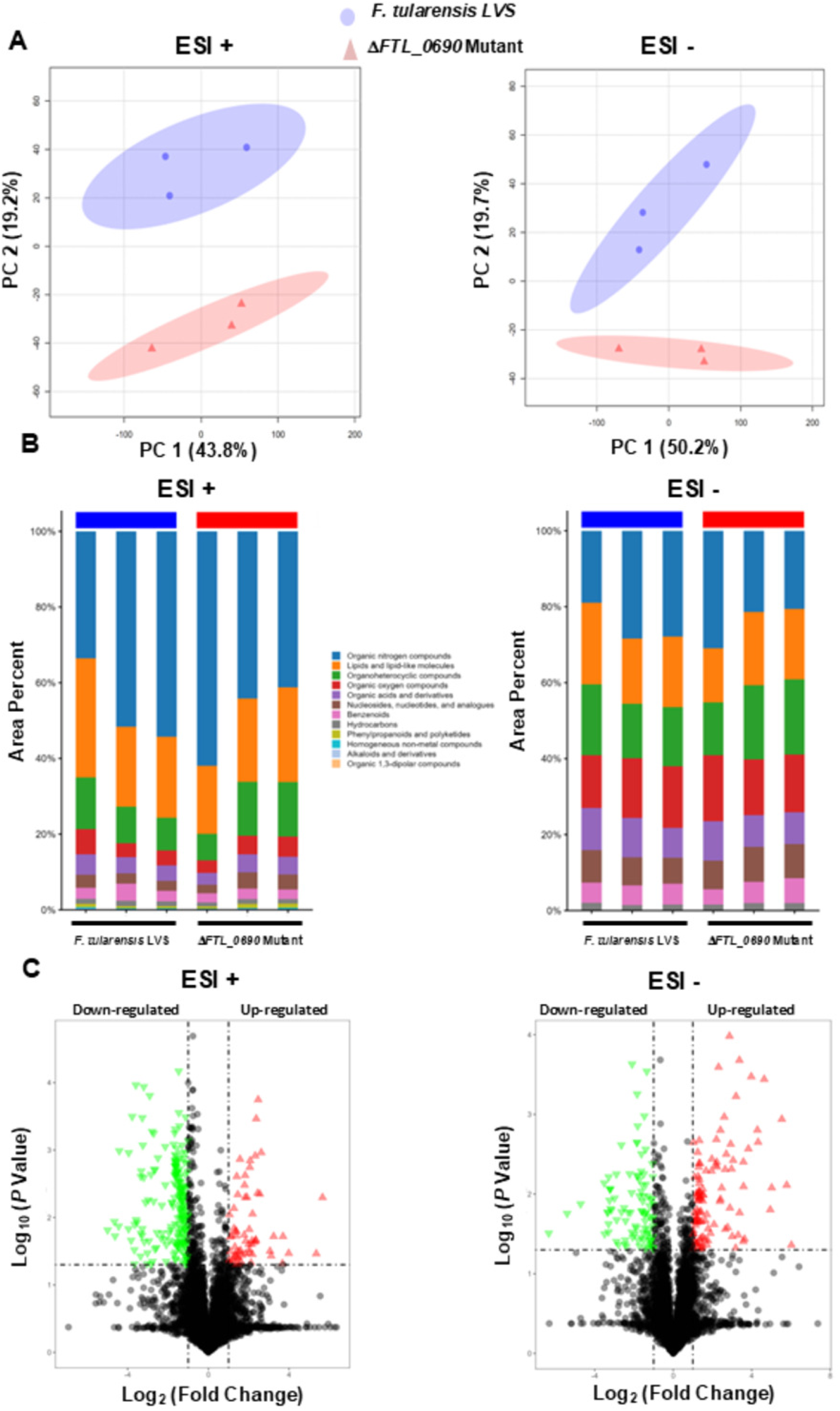
Untargeted metabolomic analysis of wild-type *Francisella tularensis* LVS and the Δ*FTL_0690* mutant. **(A)** Principal component analysis (PCA) of metabolomic profiles acquired in positive-ion (ESI+) and negative-ion (ESI−) electrospray modes. Blue circles represent wild-type *F. tularensis* LVS and red triangles represent the Δ*FTL_0690* mutant. Ellipses denote 95% confidence intervals, and the percentage of variance explained by each principal component is indicated on the axes. **(B)** Relative distribution of annotated metabolite superclasses expressed as area percent for wild-type *F. tularensis* LVS and the Δ*FTL_0690* mutant in ESI+ (left) and ESI−(right) modes. Metabolites are grouped by chemical class as indicated in the legend. **(C)** Volcano plots showing differential metabolite abundance between the Δ*FTL_0690* mutant compared to wild-type *F. tularensis* LVS in ESI+ (left) and ESI− (right) datasets. Log₂ fold change (mutant versus wild type) is plotted against −Log₁₀ (p value). Significantly up-regulated (red) and down-regulated (green) metabolites are highlighted based on the indicated fold-change and significance thresholds, with unaltered metabolites shown in gray.

Untargeted metabolomic profiling revealed clear genotype-dependent differences between wild-type *Francisella tularensis* LVS and the Δ*FTL_0690* mutant in both positive- and negative-ionization modes. In ESI+ mode, organic nitrogen compounds and lipids/lipid-like molecules dominated the metabolome in both strains; however, the Δ*FTL_0690* mutant exhibited a relative redistribution characterized by reduced organic nitrogen compounds and increased lipid-associated metabolites, along with higher contributions from organoheterocyclic and organic oxygen compounds. In ESI− mode, loss of Δ*FTL_0690* resulted in a distinct shift toward increased organic oxygen compounds, organic acids and derivatives, and phenylpropanoids/polyketides, accompanied by a relative decrease in lipid-associated metabolites. Across both ionization modes, minor metabolite classes remained low and largely unchanged. Taken together, the ESI+ and ESI−datasets demonstrate that deletion of Δ*FTL_0690* does not eliminate major metabolite classes but instead leads to a redistribution of metabolite superclasses, indicative of altered metabolic flux rather than loss of metabolic capacity (Fig. 4B). The consistent genotype-dependent differences observed across both ionization modes support a role for acyl-CoA-synthetase in maintaining metabolic homeostasis in *F. tularensis* LVS, with its absence resulting in coordinated changes affecting nitrogen-containing metabolites, lipid metabolism, and oxygenated small molecules.

Volcano plot analyses further defined the metabolic changes resulting from *FTL_0690* deletion. In both ESI+ and ESI− modes, numerous metabolites met the criteria for significant differential abundance. The Δ*FTL_0690* mutant exhibited both upregulated and downregulated metabolites relative to wild-type *F. tularensis* LVS. Notably, many of these metabolites were associated with lipid metabolism, suggesting that deletion of acyl-CoA synthetase disrupts key lipid biosynthetic or degradation pathways (Fig. 4C).

Deletion of *FTL_0690* led to broad alterations in metabolite abundance, consistent with changes in membrane composition, oxidative stress responses, and metabolic adaptation. In the ESI+ conditions, the Δ*FTL_0690* mutant exhibited increased levels of megastigmadienones, ionols, centarol, and zeranol. In parallel, fatty acid-derived metabolites were elevated in the mutant, including long-chain fatty alcohols (arachidoyl alcohol, undecanediol) and fatty amides such as N-stearoyl threonine, suggesting impaired β-oxidation and diversion of fatty acids toward membrane-associated or signaling lipids. Increased abundance of cyclic and aromatic metabolites (phenylacetone, methylindene, bicyclohexanone, and piperazine) further indicated metabolic rerouting and stress-associated overflow from disrupted central metabolism. In contrast, reduced levels of muricatacin and 3-oxo-2-cyclopentane-1-octanoic acid, lipid-derived metabolites associated with bioactive and antimicrobial functions, indicated a selective loss of lipid signaling pathways (Fig. 5A).

**FIGURE 5.**
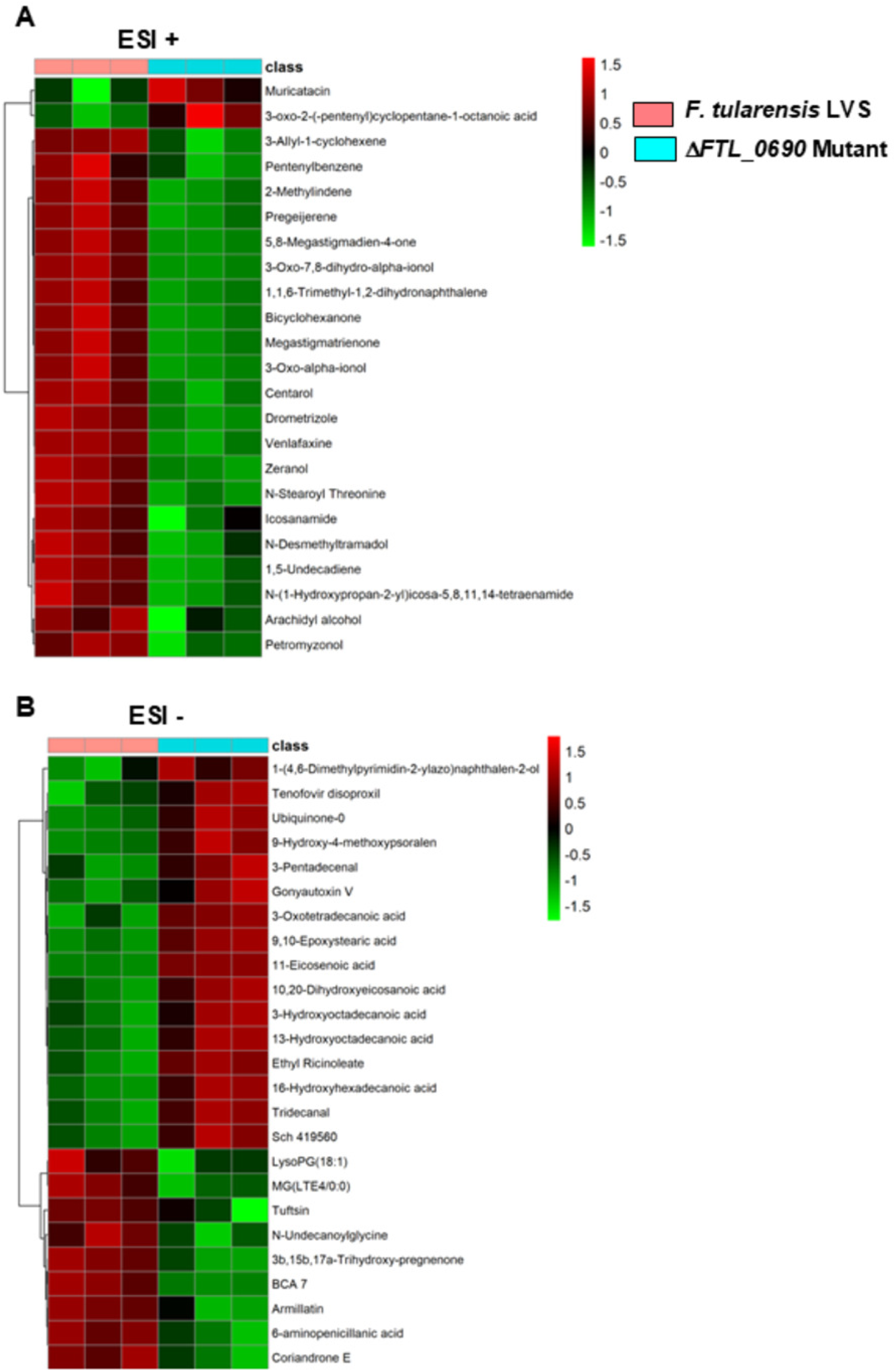
Differentially abundant metabolites in wild-type *Francisella tularensis* LVS and the ΔFTL_0690 mutant. **(A)** Heatmap of significantly altered metabolites detected in positive-ion mode (ESI+). Columns correspond to technical replicates of wild-type *F. tularensis* LVS (Pink) and the Δ*FTL_0690* mutant (Cyan), and rows represent individual annotated metabolites. **(B)** Heatmap of significantly altered metabolites detected in negative-ion mode (ESI−), displayed as in panel A. For both panels, metabolite abundances were normalized and scaled by Z-score across samples, with red indicating higher relative abundance and green indicating lower relative abundance. Metabolites were hierarchically clustered based on abundance patterns.

Under ESI− conditions, the Δ*FTL_0690* mutant displayed coordinated reductions in metabolites associated with energy metabolism, lipid processing, and stress defense relative to the wild-type *F. tularensis* LVS. Decreased ubiquinone-O level is consistent with impaired respiratory electron transport and reduced ATP synthesis. Reduced levels of β-oxidation-related metabolites (3-oxostearic and 10-oxododecanoic acids), fatty alcohols (tridecanol), and modified fatty acids (hydroxylated and epoxy species) further indicated disrupted fatty acid catabolism and diminished capacity for membrane remodeling. In addition, the lower abundance of stress- and defense-associated metabolites (gonyautoxin V and 3-hydroxy-4-methoxypsoralen), together with reduced xenobiotic-associated compounds, suggests compromised stress tolerance and detoxification capacity. The Δ*FTL_0690* mutant exhibited elevated levels of LysoPG(18:1) and an accumulation of N-undecanoylglycine, indicating increased phospholipid turnover and compensatory production of surfactant-like lipids, both of which are indicators of altered membrane integrity. Increased levels of BCA-7-related metabolites and steroid derivatives further reflected metabolic rerouting toward alternative energy sources and secondary metabolic pathways. Additionally, altered MG (LTE4/0.0) levels indicated changes in bioactive lipid signaling (Fig. 5B). In summary, the differential metabolite expression profiles observed under both ionization modes reveal that deletion of *FTL_0690* alters lipid metabolism.

### Loss of *FTL_0690* compromises membrane integrity

KEGG pathway analysis of the metabolomic data revealed significant alterations in glycerolipid and glycerophospholipid metabolism in the Δ*FTL_0690* mutant, pathways that are central to bacterial membrane structure and stability. Based on these findings, we examined whether deletion of *FTL_0690* affected membrane integrity using SDS. Wild-type *F. tularensis* LVS, the Δ*FTL_0690* mutant, and the complemented strain (Δ*FTL_0690* + p*FTL_0690*) were exposed to 1% SDS for 3 hours, followed by CFU enumeration. While all strains showed comparable growth under untreated conditions, exposure to SDS resulted in a pronounced reduction in the viability of the Δ*FTL_0690* mutant compared to wild-type *F. tularensis* LVS. In contrast, the complemented strain displayed survival comparable to that of wild-type bacteria (Fig. 6A and B). These data demonstrate that deletion of *FTL_0690* significantly increases sensitivity to SDS-mediated membrane disruption and that reintroduction of *FTL_0690* restores resistance, confirming a role for acyl-CoA synthetase in maintaining membrane integrity.

**FIGURE 6.**
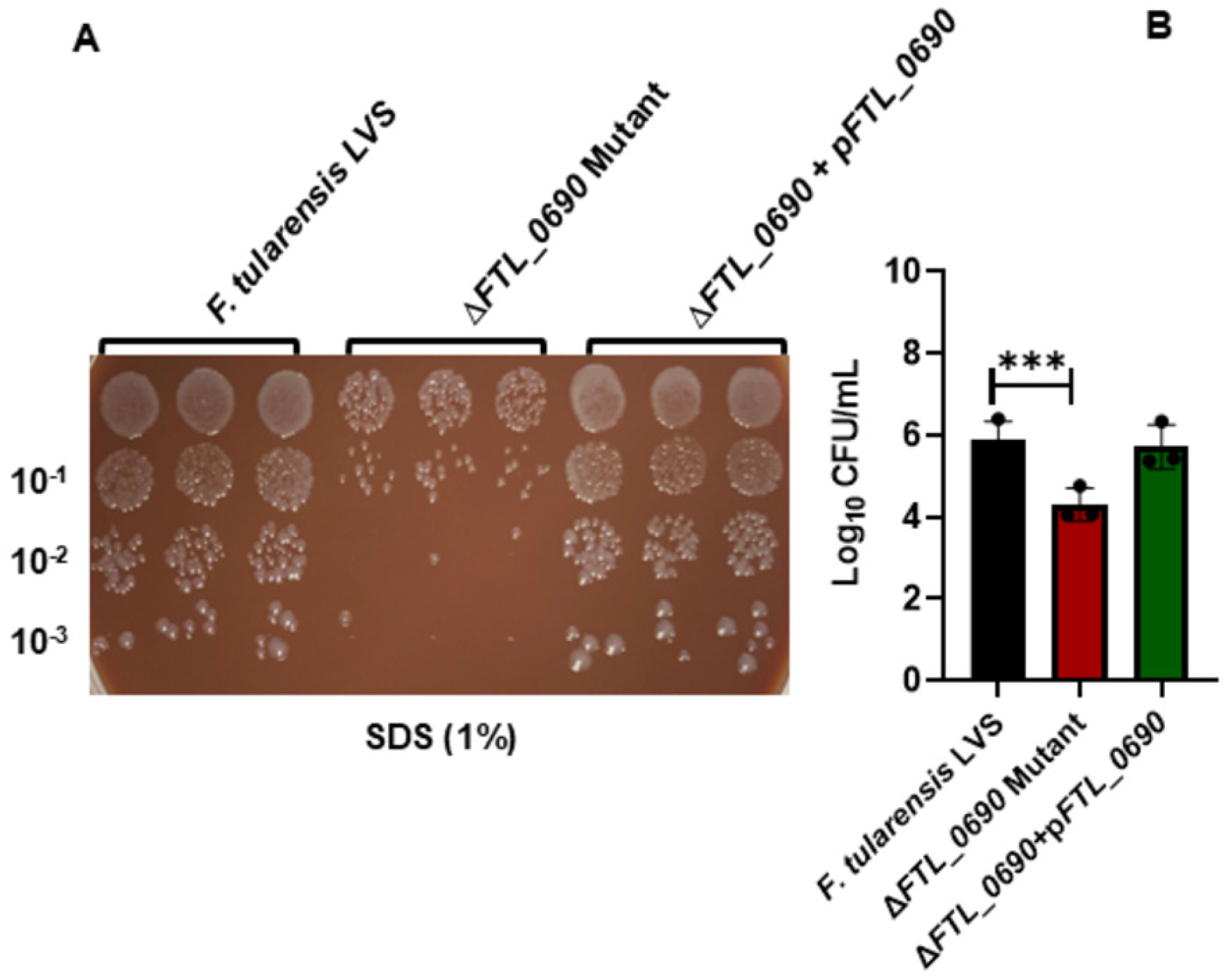
Loss of *FTL_0690* compromises detergent tolerance of *F. tularensis* LVS. Cultures of the indicated strains were grown on MH-chocolate agar plates, harvested, and resuspended in Brain-Heart Infusion broth to an OD_600_ of 0.2. Cultures were exposed to 1% sodium dodecyl sulphate (SDS) and incubated at 37°C with shaking for 3 hours. Following exposure, aliquots were spotted onto MH-chocolate agar plates to assess viability, or **(B)** were serially diluted and plated to enumerate colony-forming units (CFUs). Untreated cultures served as controls. Each spot in A represents an independent technical replicate (n = 3). Bacterial counts in B are expressed as Log₁₀ CFU/ml and represent the mean ± SEM from three experiments conducted. Statistical significance was determined using one-way ANOVA. \*\*\**P* ≤ 0.001.

### Dose-dependent virulence of the Δ*FTL_0690* mutant following intranasal infection

To assess the contribution of *FTL_0690* to virulence across a range of infectious doses, C57BL/6J mice were infected intranasally with the Δ*FTL_0690* mutant at three inoculum doses (2 × 10⁴, 2 × 10⁵, and 2 × 10⁶ CFU) and monitored for morbidity and mortality for up to 21 days. The Δ*FTL_0690* mutant exhibited a clear dose-dependent increase in virulence. At the lowest dose (2 × 10⁴ CFU), mice showed delayed onset of morbidity, and partial survival over the observation period of 21 days. Increasing the inoculum to 2 × 10⁵ CFU resulted in earlier, and 100% mortality. At the highest dose (2 × 10⁶ CFU), infection with the Δ*FTL_0690* mutant caused mortality occurring within a short timeframe post-infection (Fig. 7). Together, these data demonstrate that despite defects in oxidative stress resistance, membrane integrity, and intracellular replication observed in vitro, deletion of *FTL_0690* does not diminish the ability of *F. tularensis* to cause lethal infection in the murine intranasal infection model.

**FIGURE 7.**
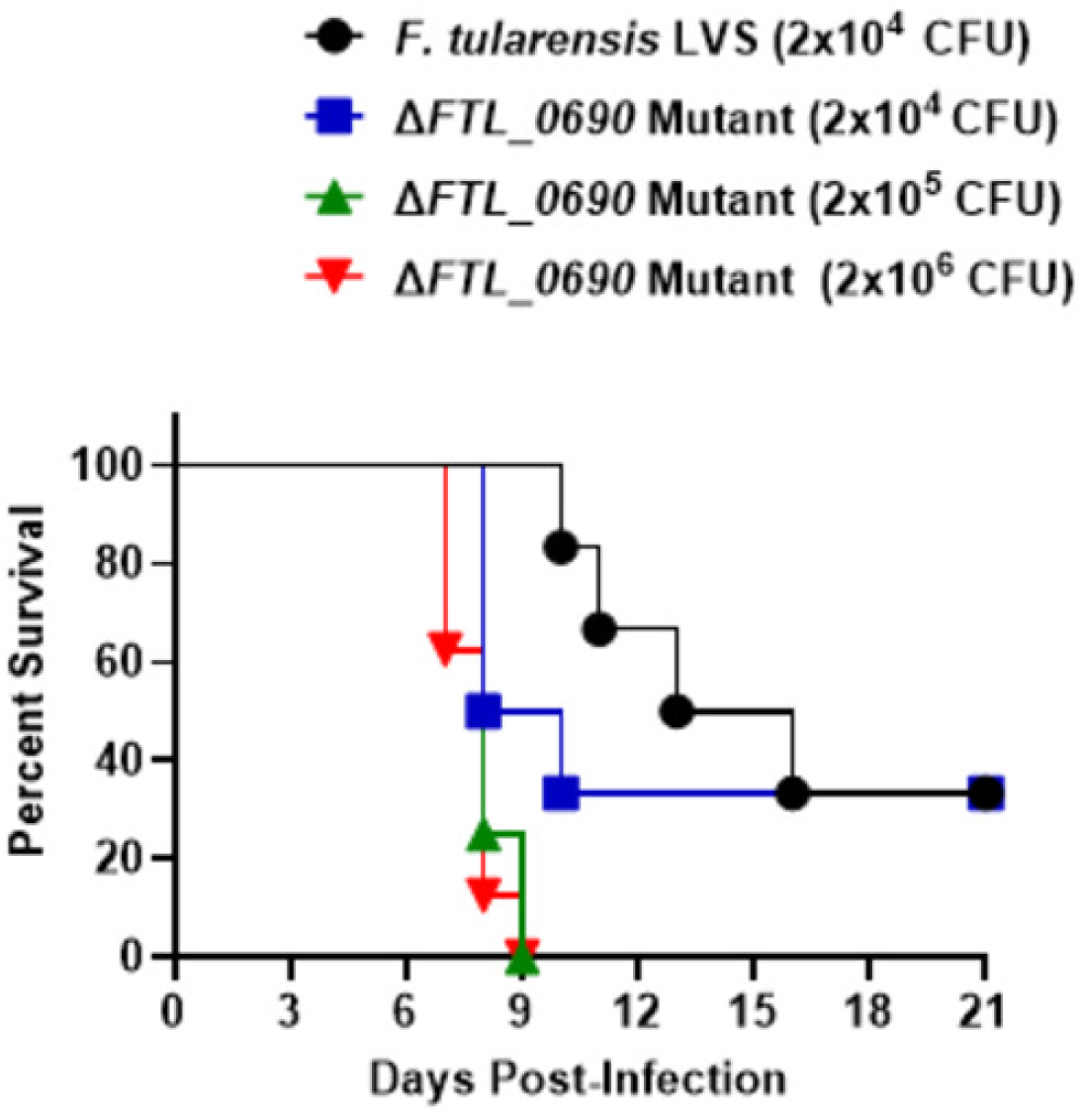
Virulence of the Δ*FTL_0690* mutant in mice. Kaplan-Meier survival curves showing percent survival over time in C57BL/6 mice (n = 8 mice per group) infected intranasally with the indicated doses of wild-type *F. tularensis* LVS or the Δ*FTL_0690* mutant and monitored daily for survival.

## Discussion

Acyl-CoA synthetases catalyze the ATP-dependent activation of fatty acids, generating acyl-CoA intermediates that integrate lipid metabolism with central carbon metabolism, membrane biogenesis, and cellular stress responses (14, 17, 19). In bacteria, controlled fatty acid activation supports not only β-oxidation and lipid biosynthesis but also maintenance of membrane integrity by limiting lipid peroxidation and facilitating membrane repair. Acyl-CoA pools further contribute to membrane-protective modifications, including cyclopropane fatty acid biosynthesis (32). These combined functions place acyl-CoA synthetases at a central junction between lipid flux and stress adaptation.

In *F. tularensis*, *FTL_0690* encodes a long-chain acyl-CoA synthetase that appears to serve as the primary enzyme responsible for fatty acid activation. Its genomic organization within a putative operon with *FTL_0691* and *FTL_0692* suggests coordinated regulation of a conserved metabolic module, an arrangement observed in other *Francisella* species and related intracellular bacteria (33). The proximity of *FTL_0690* to *orsR* (*FTL_0689*), which encodes an oxidative stress response regulator, further suggests a functional linkage between fatty acid metabolism and stress-responsive regulatory pathways (31). Consistent with this organization, disruption of *FTL_0690* resulted in multiple physiological defects. Both the transposon-insertion and deletion mutants exhibited increased sensitivity to oxidative stress, impaired replication within macrophages, altered lipid and metabolite profiles, and reduced membrane stability. These findings indicate that fatty acid activation contributes substantially to *F. tularensis* intracellular fitness and support a role for acyl-CoA synthetases in coordinating lipid homeostasis with oxidative stress tolerance.

The intracellular environment of the macrophage likely amplifies the importance of *FTL_0690*. *F. tularensis* encounters nutrient limitation, oxidative stress, and acidic pH during intracellular residence (34), conditions that increase the need for membrane repair and metabolic flexibility. The requirement for *FTL_0690* during sustained intracellular replication suggests that acyl-CoA synthetase activity becomes particularly important during later stages of intracellular growth, when energetic and biosynthetic demands increase. Comparable stage-specific requirements for fatty acid activation have been described in other intracellular pathogens, including *Legionella pneumophila*, *Brucella abortus*, and *M. tuberculosis* (16–18, 35–37). Metabolomic analysis of the Δ*FTL_0690* mutant further supports this interpretation, revealing perturbations in glycerolipid and glycerophospholipid metabolism, accumulation of long-chain and hydroxylated fatty acids, and secondary changes in organic acids and redox-associated metabolites. These coordinated pathway-level alterations are consistent with impaired fatty acid activation, altered membrane composition, and downstream effects on TCA-linked metabolism. Together, these data establish *FTL_0690* as a metabolic node linking fatty acid utilization, membrane homeostasis, and redox balance.

The increased sensitivity of the Δ*FTL_0690* mutant to SDS provides direct evidence of compromised membrane integrity, likely resulting from defective phospholipid incorporation and altered fatty acid composition. Such membrane perturbations offer an explanation for the observed sensitivity to oxidative stress and reduced intramacrophage survival, as envelope defects would increase susceptibility to host-derived reactive oxygen species and antimicrobial factors. Similar relationships between fatty acid activation, membrane integrity, and stress resistance have been reported in *P. aeruginosa* and *Salmonella* Typhimurium (38–40).

Despite these defects, the Δ*FTL_0690* mutant retained full virulence in a murine intranasal infection model. Similar discrepancies between in vitro phenotypes and in vivo virulence have been reported for other *F. tularensis* metabolic mutants (41–43), suggesting that metabolic redundancy or host-specific nutrient availability may mitigate the loss of *FTL_0690* during infection. This outcome contrasts with findings in *M. tuberculosis*, where disruption of acyl-CoA synthetase genes results in marked attenuation (44), highlighting organism-specific differences in metabolic requirements.

In summary, this study identifies FTL_0690 as a long-chain acyl-CoA synthetase that contributes to oxidative stress resistance, intracellular growth, and membrane integrity of *F. tularensis*. Although not essential for virulence in the murine model examined, FTL_0690 enhances bacterial fitness under conditions encountered during intracellular infection, emphasizing the importance of fatty acid activation in *Francisella* physiology.

## Acknowledgements

This work was supported by National Institutes of Health Grants R21AI51277 (CSB) and R15AI107698 (MM). The funders had no role in study design, data collection and analysis, decision to publish, or manuscript preparation. No financial conflicts of interest exist regarding the contents of the manuscript and its authors.

